# Application of mechanistic methods to clinical trials in multiple sclerosis: the simvastatin case

**DOI:** 10.1101/343442

**Authors:** Arman Eshaghi, Rogier A Kievit, Ferran Prados, Carole Sudre, Jennifer Nicholas, M Jorge Cardoso, Dennis Chan, Richard Nicholas, Sebastien Ourselin, John Greenwood, Alan J Thompson, Daniel C Alexander, Frederik Barkhof, Jeremy Chataway, Olga Ciccarelli

## Abstract

The analysis of clinical trials is limited to pre-specified outcomes, thereby precluding a mechanistic understanding of the treatment response. Multivariate mechanistic models can elucidate the *causal* chain of events by simultaneous analysis of multimodal data that link intermediate variables to outcomes of interest. A double-blind, randomised, controlled, phase 2 clinical trial in secondary progressive multiple sclerosis (MS-STAT, NCT00647348) demonstrated that simvastatin (80mg/day) over two years reduced the brain atrophy rate and was associated with beneficial effects on cognitive and disability outcomes. Therefore, this trial offers an opportunity to apply mechanistic models to investigate the hypothesised pathways that link simvastatin to clinical outcome measures, either directly or indirectly via changes in serum total cholesterol levels and to determine which is the more likely.

---

We re-analysed the MS-STAT trial in which 140 patients with secondary progressive multiple sclerosis were randomised (1:1) to receive placebo or simvastatin (80 mg/day). At baseline and after one and two years patients underwent brain magnetic resonance imaging; their cognitive and physical disability were assessed on the block design test and Expanded Disability Status Scale (EDSS). Serum total cholesterol levels were measured at each visit. We calculated the annual percentage change of brain volume loss using mixed-effects models. With multivariate mechanistic models and Bayesian mediation analyses, a cholesterol-dependent model was compared to a cholesterol-independent model.

As described previously, the simvastatin group showed a slower rate of brain atrophy and clinical deterioration (as reflected by both the EDSS and the block design test) and a faster decline in serum cholesterol levels (all *p* <0.05), when compared with placebo.

The cholesterol-independent model, in which simvastatin has a direct effect on the clinical outcome measures and brain atrophy, independent of its impact on lowering the serum cholesterol levels, was the more likely model. When we deconstructed the total treatment effect on EDSS and block design, into indirect effects, which were mediated by brain atrophy, and direct effects, brain atrophy was responsible for 31% of the total treatment effect on EDSS (beta=-0.037, 95% credible interval [CI]=−0.075, −0.010), and 35% of the total treatment effect on block design (beta=0.33, 95% CI=0.06, 0.72). The effect of simvastatin on both outcomes was independent of serum cholesterol levels (EDSS: beta=-0.139, 95% credible interval=-0.255,-0.025; brain atrophy: beta=0.32, 95% credible interval=0.09,0.54).

The effect of simvastatin on disability and cognitive worsening is partially mediated by brain atrophy but is independent of cholesterol reduction. Our mechanistic approach can be applied to other medications to elucidate the pathways underlying treatment effects in progressive MS.

## Introduction

Understanding the mechanisms by which a medication has beneficial effects on clinical and imaging outcomes is an unmet need in progressive multiple sclerosis (MS) research (Thompson *et al.*, 2018). The analysis of clinical trials is usually limited to pre-planned outcomes, thereby precluding an understanding of the possible mechanistic pathways by which a treatment could have an effect. Multivariate mechanistic models can elucidate the most plausible chain of events, by simultaneous analysis of multi-modal data into models that capture hypothesised pathways linking intermediate variables to outcomes of interest (Bollen and Long, 1992). They have been employed in clinical trials of Alzheimer’s disease (Douaud *et al.*, 2013), neurocognitive ageing (Kievit *et al.*, 2014), and more extensively in social sciences (Imai *et al.*, 2011).

The double-blind, randomised, controlled, phase 2 clinical trial in secondary progressive MS (MS-STAT) (Chataway *et al.*, 2014; Chan *et al.*, 2017) has shown that simvastatin reduces the whole brain atrophy rate and has positive effects on frontal lobe function and disability (Chataway *et al.*, 2014; Chan *et al.*, 2017). This trial offers the opportunity to apply multivariate mechanistic models to explain the pathways resulting in the observed simvastatin effects. An improved understanding of the pathways via which simvastatin has an effect on clinical and cognitive outcomes will stimulate further mechanistic research in secondary progressive MS, and will show that this methodology can be extended to other diseases to obtain insights into the mechanisms through which experimental therapies provide clinical benefit.

In this study, we re-analysed the MS-STAT trial data and modelled hypothesised pathways by which simvastatin affects changes in brain atrophy, clinical and cognitive outcome measures, either directly or indirectly via changes in cholesterol. We tested the following two hypotheses: (i) The reduced rate of brain atrophy development mediated the beneficial effects of simvastatin on clinical and cognitive scores; (ii) The reduction in serum cholesterol levels mediated the impact of simvastatin on physical and cognitive disability. We also investigated whether simvastatin had global, or regionally specific, impacts on brain atrophy.

## Material and Methods

### Participants

This was a *post hoc* study that included all participants of the MS-STAT trial [ClinicalTrials.gov registration number: NCT00647348] performed between 2008-2011 at three research centres and two brain imaging centres in the UK (Chataway *et al.*, 2014). MS-STAT was a phase 2 double-blind randomised controlled trial whose primary and pre-planned analyses have been reported previously (Chataway *et al.*, 2014; Chan *et al.*, 2017). Briefly, the eligibility criteria were: (i) age between 18-65 years, (ii) Expanded Disability Status Scale (EDSS) (Kurtzke, 1983) of between 4.0 and 6.5, (iii) fulfilling revised 2005 McDonald criteria (Polman *et al.*, 2005), and (iv) secondary progressive MS defined by clinically-confirmed disability worsening over the preceding two years. Patients were ineligible if they had corticosteroid treatment or relapse within three months of recruitment, or had received immunomodulatory or immunosuppressive medications within six months of recruitment. Detailed eligibility criteria are available elsewhere (Chataway *et al.*, 2014).

### Randomisation

Patients were randomised (1:1) with a central server to placebo and high-dose simvastatin (80 mg per day) groups. The randomisation software automatically minimised the following variables between placebo and treatment groups: age (<45 and >= 45 years), gender, EDSS (4-5.5, and 6.0-6.5), centre (or MRI scanner), and assessing physician. Patients, treating physicians, and outcome assessors were blind to treatment allocation. The treatment allocation was masked to the first author (AE) who performed the *post hoc* analysis. Protocol for compliance with treatment and other details are explained elsewhere (Chataway *et al.*, 2014)

### Outcomes

Patients underwent magnetic resonance imaging (MRI), clinical and cognitive assessments at baseline, after one year and two years from the study entry. This study was performed following the Declaration of Helsinki (Association, 2000) and Good Clinical Practice. Berkshire Research Ethics Committee approved the protocol. Participants gave written informed consent before screening.

### Imaging protocol

Patients were scanned at each visit (three visits in total) with 3D T1-weighted, double-echo proton density (PD) and T2-weighted MRI at two imaging centres in the UK with 1.5 Tesla and 3 Tesla scanners. Scanner and MRI protocol remained unique for each participant throughout the trial. “Scanner” was a minimisation variable (as explained above) between treatment and placebo groups. We reported acquisition protocols in **Supplementary Table 1**.

### Clinical and cognitive outcomes

Patients underwent comprehensive clinical and cognitive assessments. Here, we studied those outcomes that had shown significant (or marginally significant) changes in previous reports (Chataway *et al.*, 2014; Chan *et al.*, 2017), which were the following: the total cholesterol level, Expanded Disability Status Scale, Multiple Sclerosis Impact Scale 29v2 (total score and physical subscale)(Hobart *et al.*, 2001), Wechsler Abbreviated Test of Intelligence (WASI) Block Design test (T-score) (Wechsler, 2011), Paced-auditory serial addition test (PASAT)(Gronwall, 1977), and Frontal Assessment Battery (FAB)(Dubois *et al.*, 2000). Block Design T-score had been calculated against an age-matched reference healthy group from the test manual (Chan *et al.*, 2017).

### Image analysis

We performed image analysis based on our established pipeline for patients with MS (Eshaghi *et al.*, 2017). Our goals were to extract regional volumes, T2 lesion masks and the whole brain percentage volume change with SIENA (Smith *et al.*, 2001). Briefly, the pipeline included N4-bias field correction of T1-weighted scans to reduce intensity inhomogeneity (Tustison *et al.*, 2010), constructing a symmetric within subject template for unbiased atrophy calculation (Reuter and Fischl, 2011), rigid transformation of T1, PD, and T2 sequences to this space, automatic longitudinal lesion segmentation of visible T2 lesions with Bayesian Model Selection (BaMoS) (Sudre *et al.*, 2015; Carass *et al.*, 2017), manual editing of these lesion masks and quality assurance with 3D-Slicer, filling of hypointense lesions in T1 scans (Prados *et al.*, 2016), brain segmentation and parcellation with Geodesic Information Flows (GIF) software (Cardoso *et al.*, 2015). Technical details are explained in detail in **the Supplemental Methods**. **Supplemental Figure 1** shows the steps of this pipeline. Outputs of this pipeline were the following: (i) percentage whole brain volume change (SIENA PBVC), (ii) T2 lesion masks, and (iii) regional brain volumes according to Neuromorphometrics’ atlas, which is similar to the Desikan-Killiany-Tourville (Klein and Tourville, 2012) atlas available at http://braincolor.mindboggle.info, for each region we summed volumes of the left and right hemispheres.

## Statistical analysis

### SIENA

We used a linear regression model in which the percentage brain volume change between baseline and two-year follow-up visits was the response variable. This model included treatment allocation as the variable of interest, and the following nuisance variables: age, gender, centre, and EDSS. We calculated treatment effect defined as the adjusted difference between percentage whole brain volume change of the two treatment groups, divided by the adjusted percentage whole brain volume change in the placebo group. We set the alpha level at 0.05 for all the analyses presented in this work. We adjusted univariate analyses of regional brain volumes for multiple comparisons with the false discovery rate method in R. We used percentage brain volume changes to calculate the effect size (or Cohen’s d) between placebo and treatment groups and compared it with the original report of this trial that used a different image analysis pipeline (Cohen’s d= 0.410 (Chataway *et al.*, 2014)).

### Univariate analysis of T2 lesion load, clinical and cognitive changes

Since the focus of this study was on dynamic changes, we extended the previous analyses (Chan *et al.*, 2017) of clinical and cognitive outcomes–which were performed as pairwise average comparisons at each baseline and year two visit–to the analyses of rates of change in the two treatment and placebo groups. We aimed to identify variables with a significant difference in their *rates* of changes between the two groups including all the three visits, and to include them in multivariate mechanistic models (see below). We used univariate linear mixed-effects models in which fixed-effects were time (years from the study entry), and the interaction of time with treatment allocation. Random effects included time nested in “participant”. To allow for repeated measures, we included random intercept and slope as correlated random effects. In these models, dependent variables were cognitive or clinical outcomes (seven separate models for T2 lesion load, PASAT, block design, EDSS, Frontal Assessment Battery, and Multiple Sclerosis Impact Scale 29v2 total and its physical subscale). We included age, gender, and centre as extra (nuisance) fixed-effects variables. We used NLME package (Pinheiro *et al.*, 2017) version 3.1-131 inside R version 3.4.0 (R Core Team, 2014).

### Multivariate analysis

We performed multivariate analyses in the following steps:

i. Variable selection: to limit the analysis to measures with significant rates of change.
ii. Model construction: to formulate mechanistic models as statistical hypotheses.
iii. Model selection: to choose the most likely hypothesis.
iv. Parameter estimation: to quantify, in the most likely model, pathways between serum, imaging, cognitive, patient-reported, and clinical variables.

### Variable selection and model construction

We implemented multivariate analysis with structural equation modelling using Lavaan package version 0.5-23 (Rosseel, 2012) in R. Structural equation models allow simultaneous fitting of many regression models to quantify pathways across variables. We included outcomes from the univariate analyses (explained above) that had significant differences in their rate of change between placebo and simvastatin groups. Since nuisance variables (age, gender, and centre) did not affect the above univariate analyses, we did not include them in multivariate models. We only entered the physical subtest of Multiple Sclerosis Impact Scale-29v2 (instead of the total score) in structural equation models, because changes in this subtest drove the change in total score. We calculated the difference between baseline and second-year values for each variable and divided it by two. We refer to this as the *annualised change* throughout this manuscript.

We hypothesised two *a priori* models to explain relationships between these variables according to the literature (Bosma *et al.*, 2015; Larochelle *et al.*, 2016) and on the basis of our opinion, that are shown in **Figure 3**. The first is a cholesterol-mediated model, in which the effects of simvastatin on clinical measures (both physical and cognitive) and brain atrophy are mediated by changes in cholesterol (**Figure 3, (A)**). The second is a cholesterol-independent model, in which simvastatin has a direct effect on the clinical and MRI outcome measures, independently by its effect on serum cholesterol levels (**Figure 3, (B)**). In both models, the rate of brain atrophy development has a direct effect on clinical change, as measured by the Expanded Disability Status Scale, Block Design and the Multiple Sclerosis Impact Scale-29v2 **Figure 3**). Additionally, in both models, Multiple Sclerosis Impact Scale 29v2 is included as the last variable in the cascade of events, because it is a subjective patient-reported questionnaire expected to reflect the consequences of clinical and cognitive changes.

### Model selection and parameter estimation

We fitted both the cholesterol-mediated and cholesterol-independent model (**Figure 3**) using full-information maximum likelihood to adjust for missingness, and with the robust standard-errors to account for non-normality (e.g., EDSS). We assessed the goodness-of-fit for each model and reported the parameters for the most likely model. To evaluate overall fit of a model we used comparative fit index (CFI; compares the fit of the model with a model with uncorrelated variables; acceptable fit>0.95, good fit >0.97), standardised root mean square residual (SRMSR; square root of the average of the covariance of residuals, good fit<0.08) and root-mean-squared error of approximation (RMSEA; discrepancy between the model and population covariance; good fit <0.06) (Hu and Bentler, 1999). To estimate the relative quality of a model given the data, we calculated information criteria (Akaike information criterion [AIC], and Bayesian information criterion [BIC]) of each model. Raw AIC and BIC values do not have a meaningful scale; therefore, we calculated Akaike and Schwarz weights to represent conditional probability of each model given the data directly (Wagenmakers and Farrell, 2004). To have an unbiased estimate we calculated fit measures (mentioned above) iteratively on 1000 bootstrap samples and reported the median of bootstrap results with 95% confidence intervals.

### Bayesian post hoc mediation models

To calculate how much of the total treatment effect was mediated by intermediate variables we constructed *post hoc* models for variables involved in the significant pathways of *a priori* models (explained above). Each *post hoc* model included three variables: treatment, an intermediate variable and a final outcome. Intermediate and outcome variables were the rates of annual change of the following variables: total cholesterol level, brain atrophy, Expanded Disability Status Scale, and Block Design score. Here, we used Bayesian multivariate models to report credible intervals, especially for those of cholesterol-mediated pathways, instead of *p*-values and confidence to allow an easier interpretation of non-significant findings. This enabled testing whether the lack of significant cholesterol-mediated effects were because of lack of statistical power or there was evidence for the absence of cholesterol-mediation effects of simvastatin (HARTUNG *et al.*, 1983; Altman and Bland, 1995). We used Blavaan package version 0.3-2.283 (Merkle and Rosseel, 2015) inside R version 3.4.0 (R Core Team, 2014). We considered an effect to be significant when the 95% credible interval of a parameter did not cross zero. We discarded the first 4,000 (“burn-in” samples) and reported the next 10,000 samples as posterior distributions with Markov Chain Monte Carlo method. We used non-informative uniform priors for Bayesian analyses. These models are shown in **Figure 4** in sections (II) and (III).

### Regional rates of atrophy

We performed analyses to calculate and compare regional atrophy rates with a univariate mixed-effects model including age, gender, centre, and total intracranial volume to adjust for the head size (Malone *et al.*, 2015). We reported brain regions that had a significant rate of change in the combined treatment and placebo groups as well as separate rates for either of these groups. Further details of statistical modelling are in the **Supplemental Material**.

## Results

### Simvastatin effects on brain atrophy, clinical measures and serum cholesterol levels

Out of 140 randomised, 131 participants completed the trial and were analysed (see **Figure 1** for available data at each visit). The baseline characteristics of the participants were given in the main publication of the MS-STAT trial, and summarised in **Supplementary Table 2;** no differences in whole brain volumes, Expanded Disability Status Scale, block design T-score Multiple Sclerosis Impact Scale 29v2 and serum cholesterol levels were seen between the treated and placebo group.

**Figure 1.**
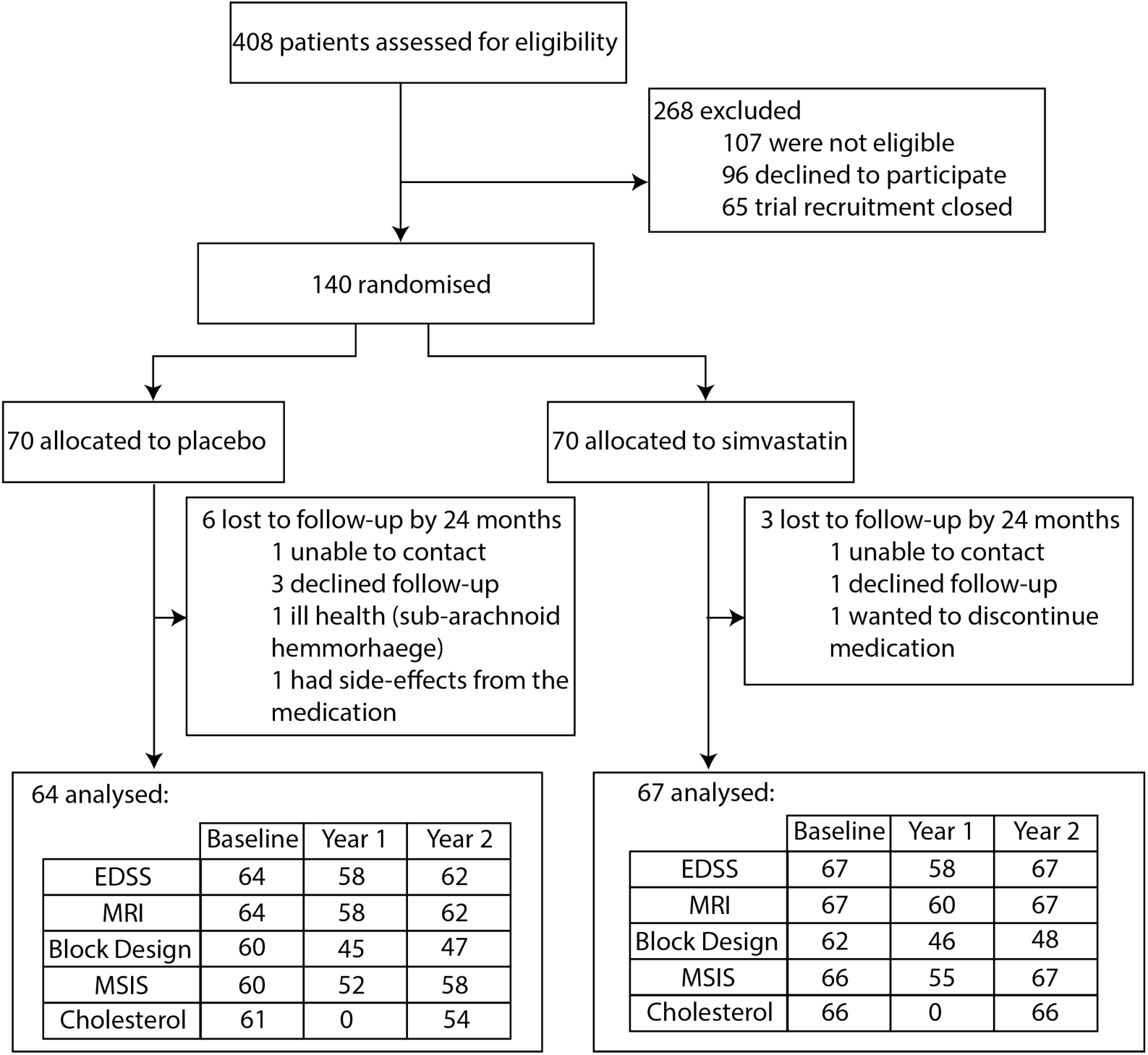
Trial profile and available data. **Trial profile and available data.** This diagram shows the flow of participants from screening to inclusion in the MS-STAT trial. Available clinical, cognitive, and imaging variables are shown in the table for all the three visits. EDSS; Expanded Disability Status Scale, MRI; magnetic resonance imaging, MSIS; Multiple Sclerosis Impact Scale.

The rate of whole brain volume loss over two years was faster in the simvastatin group than in the placebo arm: −0.42 (SD=0.50) vs. −0.657[SD=0.62), Cohen’s *d* or effect size=0.409, *p*=0.002) (**Figure 2**). The adjusted difference of the percentage brain volume change between the placebo and active treatment arm was 0.245 (95% confidence interval=0.087 to 0.403). This was similar to the original report of this trial, which used a different image analysis method (0.254, 95% confidence interval: 0.087 to 0.422). The rate of annual EDSS worsening was faster in the simvastatin group than in the placebo arm (estimated rate ± standard-error 0.08 ± 0.04 vs 0.21 ± 0.03, *p*=0.002) (**Figure 2**). Patients on simvastatin showed a significant difference in the rate of change of block design (0.92 ± 0.45 vs. −0.13 ± 0.33, *p*=0.04), and on the physical subtest of the Multiple Sclerosis Impact Scale 29v2 (0.26 ± 0.97 ± 0.72 *vs*. 2.37 ± 0.75*p*=0.03) compared to patients on placebo (**Figure 2**). There was a statistically significant decline of the total cholesterol levels in the simvastatin group compared to the placebo arm (−0.68±0.07 mmol/year *vs.* −0.01±0.05 mmol/year, *p*<0.001) (**Figure 2**). There were no differences in rates of change between treatment and placebo groups in PASAT and Frontal Assessment Battery. **Figure 2** shows the rate of change between baseline and second-year visits. There was no treatment effect on T2 lesion volume accumulation (see **Supplemental Material** for results of the lesion volume).

**Figure 2.**
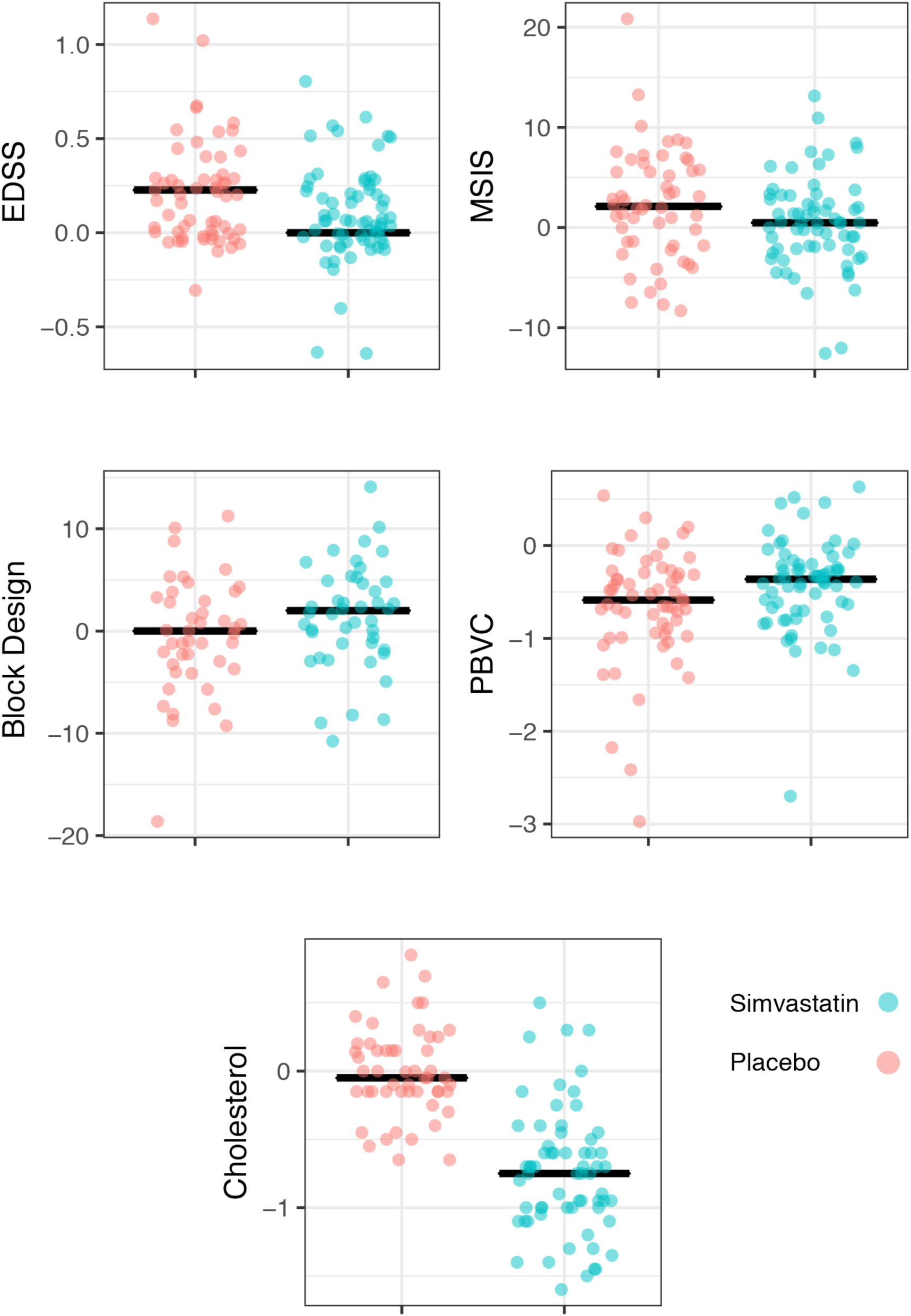
Annualised change in outcomes with a significant treatment effect. **Outcomes with a significant treatment effect**. The annual rate of changes for MRI, clinical, cognitive and patient-reported outcomes included in the mechanistic models (EDSS rates are jittered vertically by 0.1 to enable visualising overlapping values). In each of the four plots, horizontal black lines show the medians of the variable shown on *y-*axes, for placebo (blue) and statin groups (red). EDSS; Expanded Disability Status Scale, MSIS; Multiple Sclerosis Impact Scale, PBVC; percentage brain volume change.

### Effect of simvastatin on clinical outcomes is partly independent of its effect on cholesterol

The cholesterol-independent model, in which simvastatin has a direct effect on the clinical and MRI outcome measures, independently by its impact on lowering the serum cholesterol levels, was the more likely model (**Figure 3 (B**). This model showed a better overall fit than the cholesterol-mediated model. Bootstrapped fit measures for the cholesterol-independent model were the following: CFI = 0.95 (95% CI=0.86, 1), SRMR = 0.049 (95% CI= 0.02,0.07), RMSEA = 0.11 [90%CI=0, 0.18], AIC = 1800 (95% CI=1719, 1892), BIC = 1860 (95% CI=1779,1952), Akaike weight=0.71, Schwarz weight = 0.46). This means that cholesterol-independent model was 42.24 times 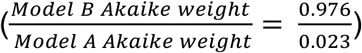 more likely than the cholesterol-mediated model regarding Kullback–Leibler discrepancy. Cholesterol-independent model was 2.38 times 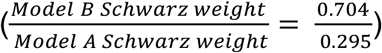 more likely regarding Schwarz weights than the cholesterol-mediated model. **Figure 3** shows fit measures for both models.

**Figure 3.**
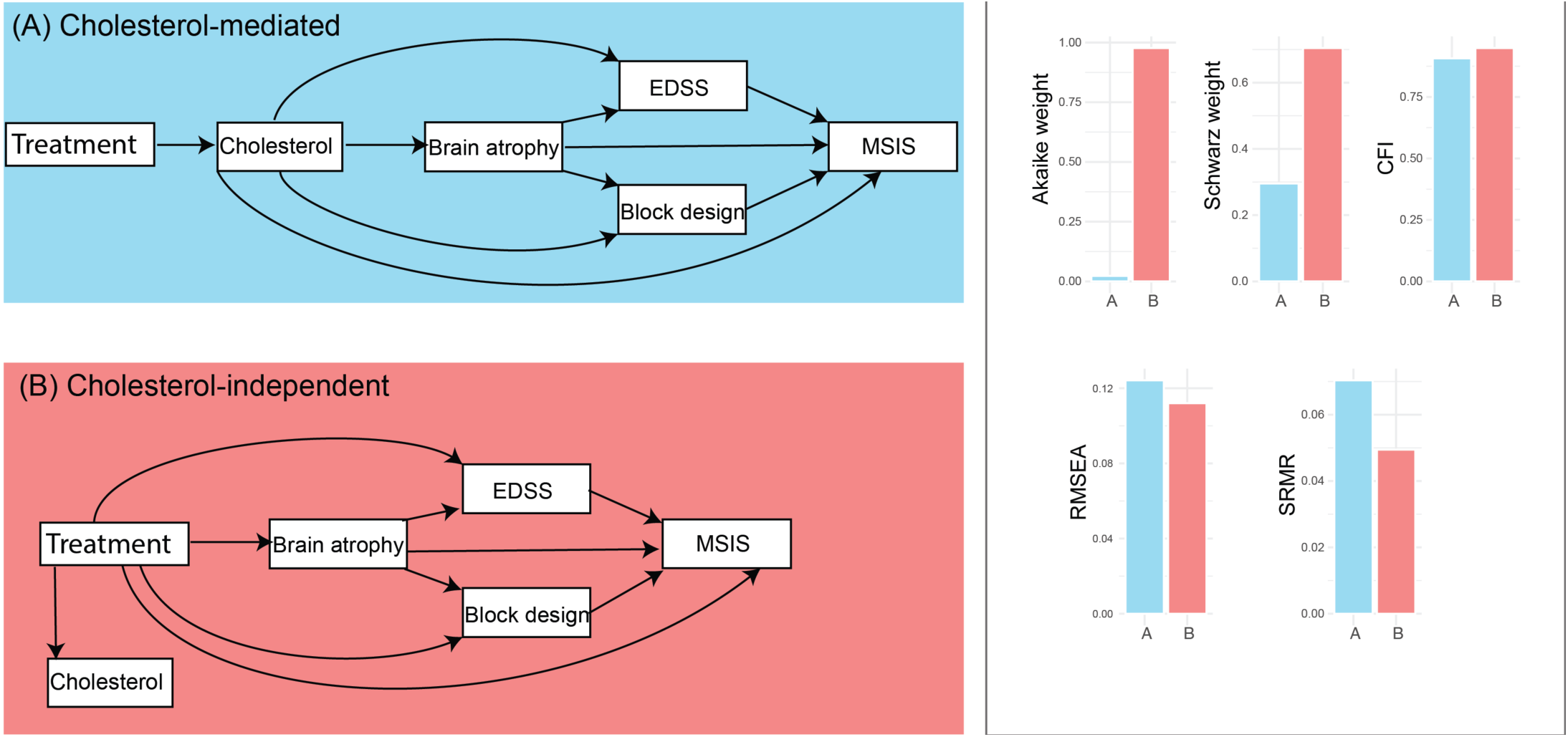
Candidate *a priori* models that that explain the causal chain of event that leads to simvastatin effects on clinical scores. **Cholesterol-mediated and cholesterol-indipendent models that explains the effect of simvastatin on clinical scores.** Model (A) or cholesterol-mediated model assumes that the cholesterol-lowering effect of simvastatin is the cause of slowing of the brain atrophy and disability worsening. Model (B) or cholesterol-independent (or pleiotropic) model assumes that the cholesterol-lowering effect of simvastatin is independent of its effect on brain atrophy and clinical outcomes. In both models, a lower rate of brain atrophy development has an effect on the clinical change, as measured by the Expanded Disability Status Scale, Block Design and the Multiple Sclerosis Impact Scale-29v2. Additionally, in both models, Multiple Sclerosis Impact Scale 29v2 is included as the last variable in the cascade of events, because it is a subjective patient-reported outcome measure. All the variables are “annualised”, which represent annual rates of change between baseline and second-year follow-up visits. Each rectangle represents a variable. Arrows represent multivariate regressions, where an arrow starts from a predictor and points to the dependent variable. The bar plots in the right column compare fit-measures that are shown on the y-axis of each of the five bar plots with models (A) and (B) on the x-axis. Blue corresponds to cholesterol-mediated model and red to cholesterol-independent model. Fit measures suggest that cholesterol-independent model (or model B) was the most likely model given data, because it had a higher Akaike and Schwarz weights, higher CFI, lower SRMR, and lower RMSEA. EDSS; Expanded Disability Status Scale, PBVC; percentage brain volume change, MSIS; Multiple Sclerosis Impact Scale. CFI; confirmatory factor index, SRMR; standardised root mean square residual, RMSEA; root mean squared error of approximation.

When we decomposed the total treatment effect on EDSS and block design, into indirect effects, which were mediated by brain atrophy, and direct effects, brain atrophy mediated 31% of the total treatment effect on EDSS (beta=-0.0364 95% credible interval [CI]=-0.075, −0.010), and 35% of the total treatment effect on block design (beta=0.324 95% CI=0.06, 0.72) (**Figure 4**). Simvastatin appeared to have direct effects on EDSS (beta=-0.164, *p*=0.047), and brain atrophy (beta=-0.364, p<0.001), but not on block design and MSIS (**Figure 4**).

**Figure 4.**
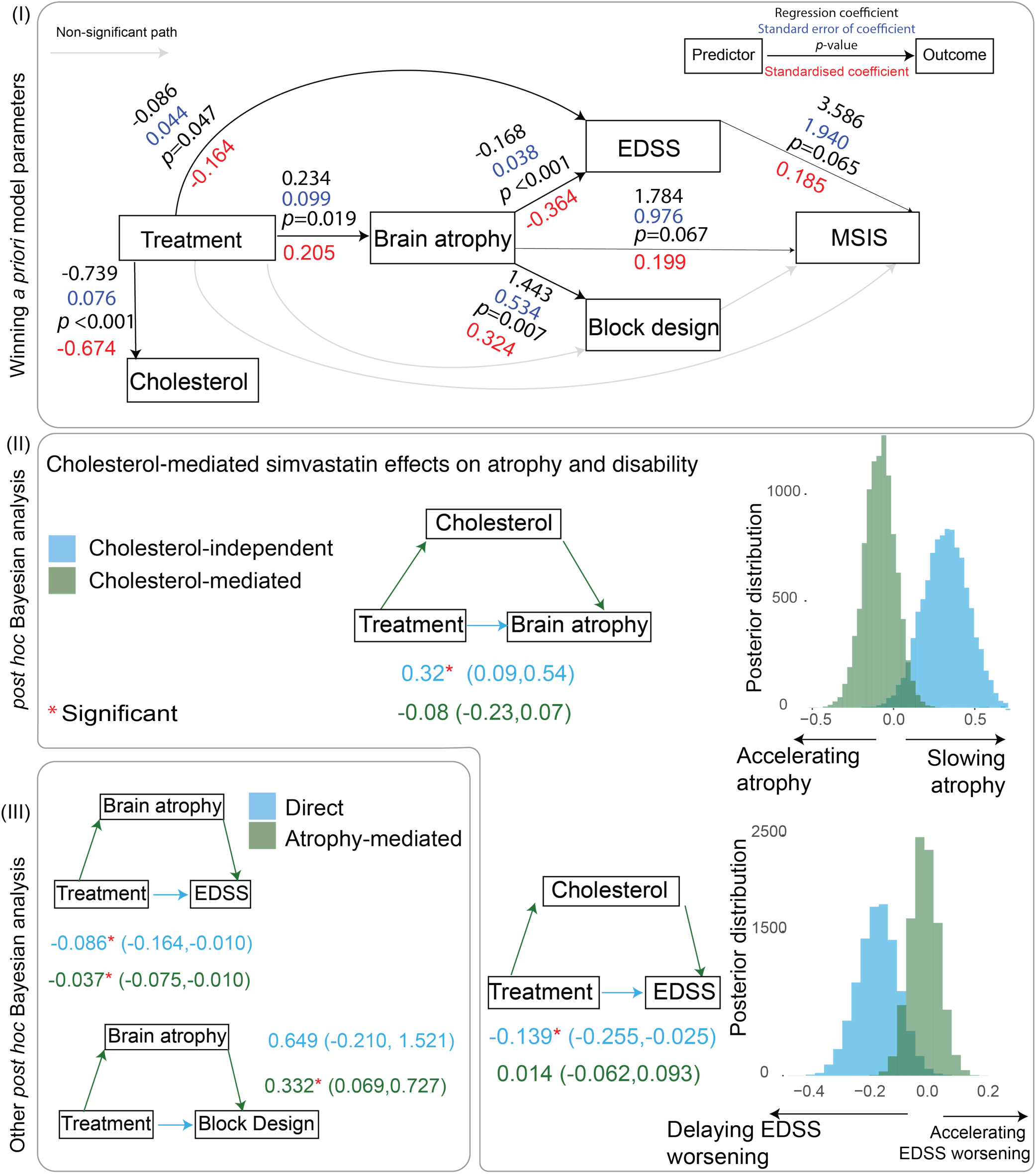
Parameter estimates of the winning model (cholesterol-independent) and *post hoc* analyses of cholesterol pathways. **Parameter estimates of the most likely model.** The section (I) shows the parameter estimates of the winning model (model (B) in **Figure 3**). Each arrow is a regression “path” where the arrow starts from the predictor(s) and points to the dependent variable(s). Significant paths (p<0.05) are shown with bold arrows, while non-significant paths are thinner. Black numbers on each arrow represent regression coefficients and their *p*-values. Blue numbers represent standard errors of the coefficients. The red numbers represent standardised coefficients. Section (II) shows the Bayesian *post hoc* analysis of cholesterol-mediated pathway *vs* direct pathway that does not depend on cholesterol to slow brain atrophy. The results confirm that a direct pathway (cholesterol-independent) slows brain atrophy. The numbers on the left side of the section (II) show median of the posterior distribution of the model parameters, and the numbers inside parenthesis show 95% credible intervals. The 95% credible intervals of coefficients of direct pathway and cholesterol mediated pathways do not overlap, this suggests that the lack of significance in cholesterol-mediated pathway is unlikely to be due to a lack of statistical power. We used a Bayesian method to ease the interpretation of non-significant findings and to report credible intervals (rather than the confidence intervals). The section (II) also shows Bayesian mediation analyses for EDSS. The direct effect is shown in blue and the mediation effect (or indirect effect) is shown in green. The treatment effect on EDSS is, at least partly, independent from its effect on cholesterol because the 95% credible intervals do not overlap except for the range between −0.062 to −0.025. Brain atrophy mediates 31% of the treatment effect on EDSS. Section (III) shows mediation analysis for other variables. PBVC; percentage brain volume change, EDSS; Expanded Disability Status Scale, MSIS; Multiple Sclerosis Impact Scale (physical subtest).

### Post-hoc Bayesian mediation analyses confirmed that atrophy, but not cholesterol, mediated simvastatin effects on EDSS

There was no significant mediation of the treatment effect via cholesterol on atrophy (treatment➔cholesterol➔atrophy, beta=-0.08, 95% credible interval=-0.23, 0.07, **Figure 4[II]**), while in the same model there was a significant direct effect of treatment that delayed atrophy (treatment➔atrophy, beta=0.32, 95% credible interval=0.09, 0.54). Since the 95% credible intervals of these two parameters do not overlap, this suggests that the lack of statistical significance for cholesterol-mediated slowing of atrophy is not due to lack of statistical power (see **Figure 4[II]**). Thus, the treatment effect on brain atrophy is independent from its effect on cholesterol. In another mediation model, cholesterol did not have a significant treatment mediation effect on EDSS (treatment➔cholesterol➔EDSS, beta=0.014, 95% credible interval=- 0.062,0.093), while in the same model there was a significant cholesterol-independent effect (beta=-0.139, 95% credible interval=-0.255,-0.025), suggesting that treatment effect on EDSS is, at least partly, independent of cholesterol. Brain atrophy significantly (beta=-0.037, 95% credible interval=-0.075,-0.010, **Figure 4[II]**) mediated 31% of the total treatment effect on EDSS (treatment➔atrophy➔EDSS) and the remaining 69% was direct. In the mediation model with brain atrophy and block design (**Figure 4[III]**), brain atrophy mediated 35% of the total treatment effect on block design (treatment➔atrophy➔block design, beta=0.33, 95% credible interval=0.06,0.72). There were no mediation effects of EDSS or brain atrophy on MSIS (treatment➔atrophy➔MSIS or treatment➔EDSS➔MSIS, not shown in **Figure 4**).

### Regional analysis

In the analysis of the merged treatment and placebo groups several regions showed significant rate of loss over time, the fastest of which was the lateral ventricle (1.95% annual expansion [95% confidence interval: 1.53%, 2.38%]), and then the transverse temporal gyrus (estimated annual rate= −1.17% [95% confidence interval: −0.88%, - 1.46%]. Rates of volume loss in the postcentral and precentral gyri, frontal regions, anterior and middle parts of the cingulate cortex, precuneus, and the thalamus were also significant (which implies ongoing volume loss, see **Figure 5** for the full list). When comparing placebo and simvastatin groups, the rates of atrophy were numerically slower in several regions in the simvastatin group (see **Figure 5**). Only the transverse temporal gyrus showed a significant difference (*p*=0.002) in rates of change (estimated annual rate [95% confidence interval] in placebo group = −1.58% [95% confidence interval: −1.17%, −1.98%]), simvastatin group = −0.79% [95% confidence interval: - 0.22%, −1.35%]) (50% treatment effect). The spatial pattern of focal volume loss was similar between the placebo and simvastatin groups on visual inspection and qualitative comparison.

**Figure 5.**
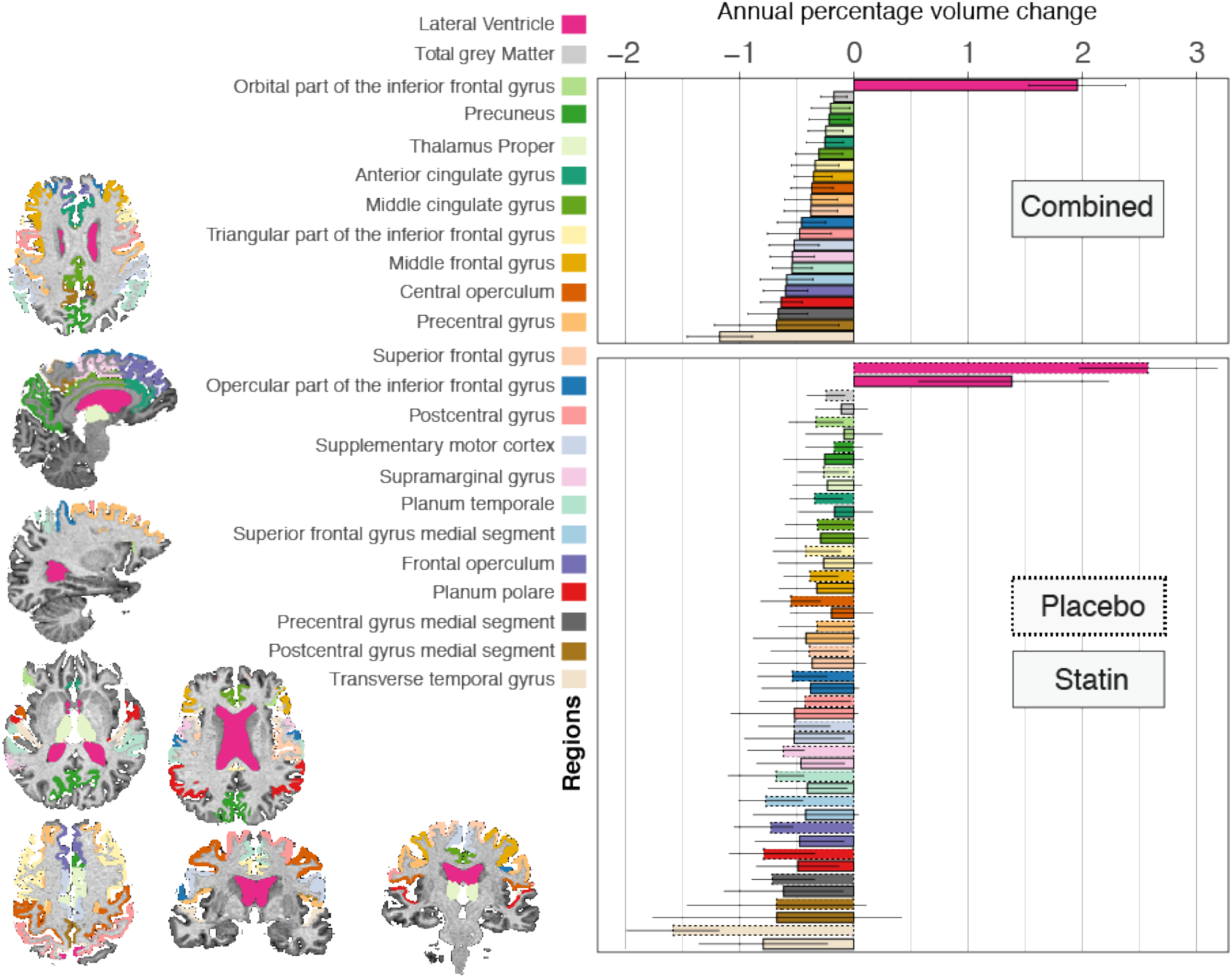
Atrophy rates in areas with a significant ongoing change. **Atrophy rates in areas with significant ongoing change.** This graph shows the adjusted annual rates of volume loss (or expansion for the lateral ventricles) which are calculated from the coefficient of the interaction of time and treatment group in the mixed-effects models constructed separately for each region. Only regions with significant volume change in the combined placebo and treatment analysis are shown (adjusted for multiple comparisons with the false-discovery method). Different colours correspond to different regions that are shown with the same appearance in left on the T1-weighted scan of one of the patients (chosen at random) and, in the right, as bar plots. Two bar plots are shown; the above shows the rate of change in the combined analysis of placebo and treatment groups on the horizontal axis and different regions on the vertical axis. The lower bar plot shows the rate of change for the same areas for placebo and simvastatin groups separately. This bar plot shows that only the transverse temporal gyrus shows a significant difference in the rate of change when comparing simvastatin and placebo groups. The error bars indicate 95% confidence interval of the rate of change.

## Discussion

In this study, we compared mechanistic hypotheses on *how* a potential neuroprotective drug, simvastatin, can influence imaging, clinical, cognitive, and patient-reported outcomes through changes in peripheral cholesterol level. The more likely model suggested that delaying of the EDSS worsening was caused by two possible effects; one by slowing of the atrophy, and another by a direct effect from simvastatin. This model also suggests that simvastatin delayed the brain atrophy and EDSS worsening independent of its cholesterol-lowering effects. Further mediation analysis with Bayesian models confirmed these findings: slowing of atrophy rate and disability worsening were more likely to be cholesterol-independent, and the slowing of disability was caused by the slowing of atrophy by simvastatin. Additionally, slowing of the atrophy also delayed the worsening of block design scores. Our work showcases the promise of mechanistic methods to compare underlying mechanisms of drug actions that are observed as treatment effects in clinical trials of progressive MS.

Cholesterol-independent model and *post hoc* mediation models suggested that a reduction in the rate of EDSS worsening was partly (31%) explained by the treatment effects on brain atrophy, and partly (69%) by a separate *direct* treatment effect. All of these effects were, at least partly, independent of the change in cholesterol levels. Our mechanistic approach, also known as mediation analysis, goes beyond correlation analysis and provides causal evidence of association between two variables. This starts by mathematically deconstructing simvastatin effects as cholesterol-mediated or cholesterol-independent and allows an indirect understanding of whether beneficial simvastatin effects are mediated via mevalonate pathway (that produces cholesterol) or not. Cholesterol is only one of the products of the 3-hydroxy-3-methyl-glutaryl-coenzyme A (HMG-CoA) reductase (part of mevalonate pathway), an enzyme that is inhibited by simvastatin. Cholesterol-independent (or pleiotropic) products of this pathway include isoprenoids that prenylate a variety of key signalling proteins that regulate cell function (Greenwood and Mason, 2007). Moreover, statins have direct effects on leukocyte adhesion that are independent of mevalonate pathway and its metabolites (Weitz-Schmidt *et al.*, 2001). Previous report of MS-STAT trial (Chataway *et al.*, 2014) demonstrated no significant effect of simvastatin on five immunological markers (IFN-γ, IL-4, IL-10, IL-17, and CD4 Fox P3). Cholesterol-independent simvastatin effects, however, may be mediated by many other immunomodulatory factors that affect leukocyte migration, antigen presentation, vasculoprotection, super oxides, and diffuse inflammation (Greenwood *et al.*, 2006; Greenwood and Mason, 2007). Taken together, our results underline the potential importance of simvastatin effects independent of lipid lowering and related pathways that do not affect HMGCoA reductase (Weitz-Schmidt *et al.*, 2001). Our results provide novel insights and stimulates further mechanistic research for drug discovery in secondary progressive MS.

The simvastatin effect on brain volume loss was driven by a general reduction in volume loss in multiple regions, but a significant effect was only seen in a region with the highest rate in the grey matter (the transverse temporal gyrus). This can be explained by a diffuse neuroprotective effect. Another novelty of our study, is that the spatiotemporal pattern of *ongoing* atrophy in patients with secondary progressive multiple sclerosis with very long disease duration (21 years), to the best of our knowledge, has never been investigated before. Our regional analysis showed that localised atrophy in the temporal lobe, frontal lobe, limbic cortex, and the basal ganglia continues relentlessly, but the pattern of ongoing atrophy was generalised as opposed to a regional loss (e.g., thalamus) (Eshaghi *et al.*, 2018). Regional susceptibility of neuroanatomical areas to neurodegeneration manifests by faster *percentage* of atrophy rates than that of the entire brain. For example, annual percentage volume loss can be up to 4% in the hippocampus in Alzheimer’s disease (Henneman *et al.*, 2009; Josephs *et al.*, 2017), while it is up to 1% for the entire brain. In MS, the deep grey matter atrophy rates can be up to 1.5% (Eshaghi *et al.*, 2018), while the whole brain atrophy is 0.6%. In this study, we found that the highest rate of loss was in the lateral ventricle–a non-specific generalised measure of atrophy. Unlike patients with early secondary progressive or primary progressive MS, none of the deep grey matter nuclei showed a higher rate than total brain rate (the thalamic atrophy rate was 0.24%), while the whole brain volume loss on average was similar to previous studies (0.65%). The slower than expected rate of atrophy in the deep grey matter suggests a floor-effect at which the decline of these structure may slow down. Our results are in line with pathological observations that generalised neurodegeneration may dominate long-standing secondary progressive MS (Frischer *et al.*, 2009; Hawker *et al.*, 2009; Carassiti *et al.*, 2017), while a more selective pattern is seen in earlier MS alongside focal inflammation that responds to immunomodulation (Frischer *et al.*, 2009; Montalban *et al.*, 2017). Although there was a general reduction in several regions in the simvastatin group, only the treatment effect on the transverse temporal gyrus was significant, which also had the highest rate of volume loss in the grey matter. Therefore, a general effect of slowing atrophy rate became detectable in a region with a higher rate. We can speculate that transverse temporal gyrus is spared until later stages of secondary progressive MS, and showed a higher rate after exhaustion of other areas.

A major difference between our study and the previous analyses of MS-STAT (Chataway *et al.*, 2014; Chan *et al.*, 2017), is that we calculated *rates* of change in imaging and clinical outcomes, rather than average differences between treatment groups at each visit which have been reported before (Chataway *et al.*, 2014; Chan *et al.*, 2017). As Chan et al. reported before, there was a significantly better frontal lobe function (as assessed by Frontal Assessment Battery) in the simvastatin group as compared to the placebo group at 24 months. However, this previous report was only limited to pre-planned statistical analysis of this trial (Chan *et al.*, 2017), and did not look at the rate of change. In this study, we performed an independent image analysis and looked at the rate of change, using all three visits (baseline, year 1 and year 2), and found that the rate was significant for the block design but not for the Frontal Assessment Battery. This is because frontal assessment battery, unlike block design, showed a ceiling effect after the first year (results are not shown) of this trial, which reduces the rate of change. We, therefore, only included block design scores in the multivariate mechanistic models. Block design evaluates the visuospatial memory and depends on fine motor coordination (as it is timed) (Groth-Marnat and Teal, 2000). While there was an association between the rate of brain volume loss and block design test, evidence for an indirect treatment effect on this cognitive outcome was weaker than Expanded Disability Status Scale. Our results demonstrate that mechanistic multivariate models can quantify and elucidate interrelations of multi-modal measures in a clinical trial.

We used a novel image analysis pipeline alongside SIENA and reproduced the original findings of the MS-STAT trial independently, which was conducted by boundary-shift integral (BSI) and different segmentation and registration methods. The differences between rates of atrophy between placebo and treatment groups were in excellent agreement (average [95%CI] difference between groups in our study: 0.245 [0.087 to 0.403], and in the original report (Chataway *et al.*, 2014): 0.254 [0.087 to 0.422]). The effect size in this study (0.409) was similar to the original report (0.410), which confirms a small to medium effect of simvastatin on brain atrophy. However, rates of percentage brain volume were slightly higher in our study. For example this rate for the placebo group was 0.587% annual loss in the original report but it was 0.657% in this study. This is a methodological artefact due to a slightly faster average atrophy rates calculated by SIENA (compared to BSI used in the original report). A previous methodological comparison showed that SIENA produces 20% faster atrophy rates, while these two methods had an excellent agreement otherwise (Smith *et al.*, 2007), which is confirmed by the adjusted difference and similar effect sizes.

It is important to note that our study is limited by its *post hoc* nature. While pre-planned statistical analyses of clinical trials are the gold-standard to compare treatments, *post hoc* analyses may nevertheless provide information to generate new hypotheses from the wealth of information collected as part of a trial.

We showed that mechanistic modelling has the potential to compare the effects of neuroprotective drugs on pathways underlying disability. Beneficial effects of simvastatin in secondary progressive MS are mainly due to the pleiotropic effects, rather than its lipid-lowering effects. Simvastatin mainly affects motor functioning directly, and indirectly by slowing atrophy rates. A weaker simvastatin effect on visuospatial memory may also exist that is mediated by slowing atrophy rates. Our approach can be extended to trials of neurodegenerative disorders to elucidate and quantify the pathways underlying disease worsening and treatment effects.

## Acknowledgement

Arman Eshaghi has received McDonald Fellowship from Multiple Sclerosis International Federation (http://www.msif.org) for this work. O. Ciccarelli, A. Thompson, and F. Barkhof have received funding from the National Institute for Health Research (NIHR) University College London Hospitals (UCLH) Biomedical Research Centre (BRC) for this work. D. Alexander has received funding for this work from EPSRC (M020533, M006093, J020990) as well as the *European Union’s Horizon 2020 research and innovation programme* under grant agreement Nos 634541 and 666992. We gratefully acknowledge Prof Chris Frost’s comments on the draft of this manuscript.

## Supplemental material

## Supplemental methods

### Image analysis

#### *1.* N4-bias field correction

We used ANTs software version 2.2(Tustison *et al.*, 2010) to correct for the scanner-field inhomogeneity in T1 scans. We used Montreal Neurological Institute intracranial mask (Boyes *et al.*, 2008) transferred with diffeomorphic registration(Avants *et al.*, 2008) to the native space to limit the correction to the cranium.

#### *2.* Symmetric within-subject template construction

We constructed an isotropic symmetric template per subject using available time-points with iterative rigid registration(Reuter and Fischl, 2011; Leung *et al.*, 2012). This step is necessary to avoid bias towards a time-point (e.g., baseline) since it distributes interpolation and segmentation errors across time-points for an unbiased atrophy calculation(Fox *et al.*, 2011).

#### 3. Symmetric transformation

We transferred T1, PD and T2 scans to the within-subject template by applying the symmetric transformation matrix. We reconstructed scans with B-spline interpolation to minimise blurring artefacts.

#### *4.* Automatic lesion segmentation

We used Bayesian Model Selection (BaMoS) to segment white matter lesions longitudinally and produce lesion masks(Sudre *et al.*, 2015, 2017; Carass *et al.*, 2017). BaMoS is a multimodal method that integrates PD, T2, and T1 segmentations to provide lesion masks.

#### 5. Manual editing

We used 3D-Slicer (https://www.slicer.org) version 4.6 to manually edit lesion masks acquired from BaMoS.

#### *6.* White matter segmentation

We used Geodesic Information Flows (GIF)(Cardoso *et al.*, 2015) version 3.0 to segment T1 scans and calculate (normal-appearing) white matter masks. This mask enables filling hypointense white matter lesions while avoiding any change in ventricular sizes(Prados *et al.*, 2016).

#### *7.* T1 hypointense lesion filling

We used a longitudinal patch-based method to fill hypointense lesions on T1 scans(Prados *et al.*, 2016). We used white matter mask from the previous step as a reference to fill hypointense lesions. This step minimises erroneous segmentation of hypointense-lesions as grey matter and increases the precision of atrophy estimates as explained elsewhere(Prados *et al.*, 2016).

#### *8.* Brain segmentation and parcellation

We used GIF to segment lesion-filled T1 scans into grey matter, white matter, and CSF and to parcellate the brain to ~120 regions according to the Desikan-KillianyTourville protocol (http://braincolor.mindboggle.info/index.html)(Klein and Tourville, 2012). We calculated the volume of each parcellated region by multiplying segmentation probability maps with the voxel volume.

To calculate whole brain percentage atrophy we used SIENA (part of FSL version 5.0)(Smith *et al.*, 2001). SIENA estimates the rate of atrophy by measuring the shift of brain edge over two separate time-points. To have consistent results between regional and global atrophy that were not limited by the differences in segmentation methods, we used GIF masks within SIENA instead of BET(Smith, 2002) and FAST(Zhang *et al.*, 2001).

### Statistical analysis

#### Regional analysis

To explore regional treatment effects, and primary drivers of the total brain atrophy we summed respective regions from left and right hemispheres and constructed linear mixed-effects models for each area (~60 models), where the volume of a given area was the dependent variable. Independent variables (fixed effects and random effects) were similar to the models used for cognitive and clinical outcomes with an additional variable for total intracranial volume to adjust for the head size(Malone *et al.*, 2015) and scanner (1.5 Tesla or 3 Tesla). First, we extracted rates of atrophy for those regions that had a significant rate of change (significant slope), after adjustment for multiple comparisons with the false-discovery rate (Benjamini and Hochberg, 1995). With a similar model, we calculated the rate of change for treatment and placebo groups.

#### There was no treatment effect on the rate of change in T2 lesion volume

At baseline, lesion volume in the placebo group was 22.14 mL (95%CI: 18.82 to 25.46), which was not different (p=0.33) from the treatment group (average=19.3, 95% CI: 13.48 to 25.12). Lesion volumes significantly increased in each group: average [95%CI] for the treatment group was 0.55 ml/year [0.25 to 0.85], and the average for the placebo group was 0.72 ml/year [0.55 to 0.87]). However, rates of change were similar between treatment and placebo groups.

## Supplemental Figures

**Supplemental Figure 1.**
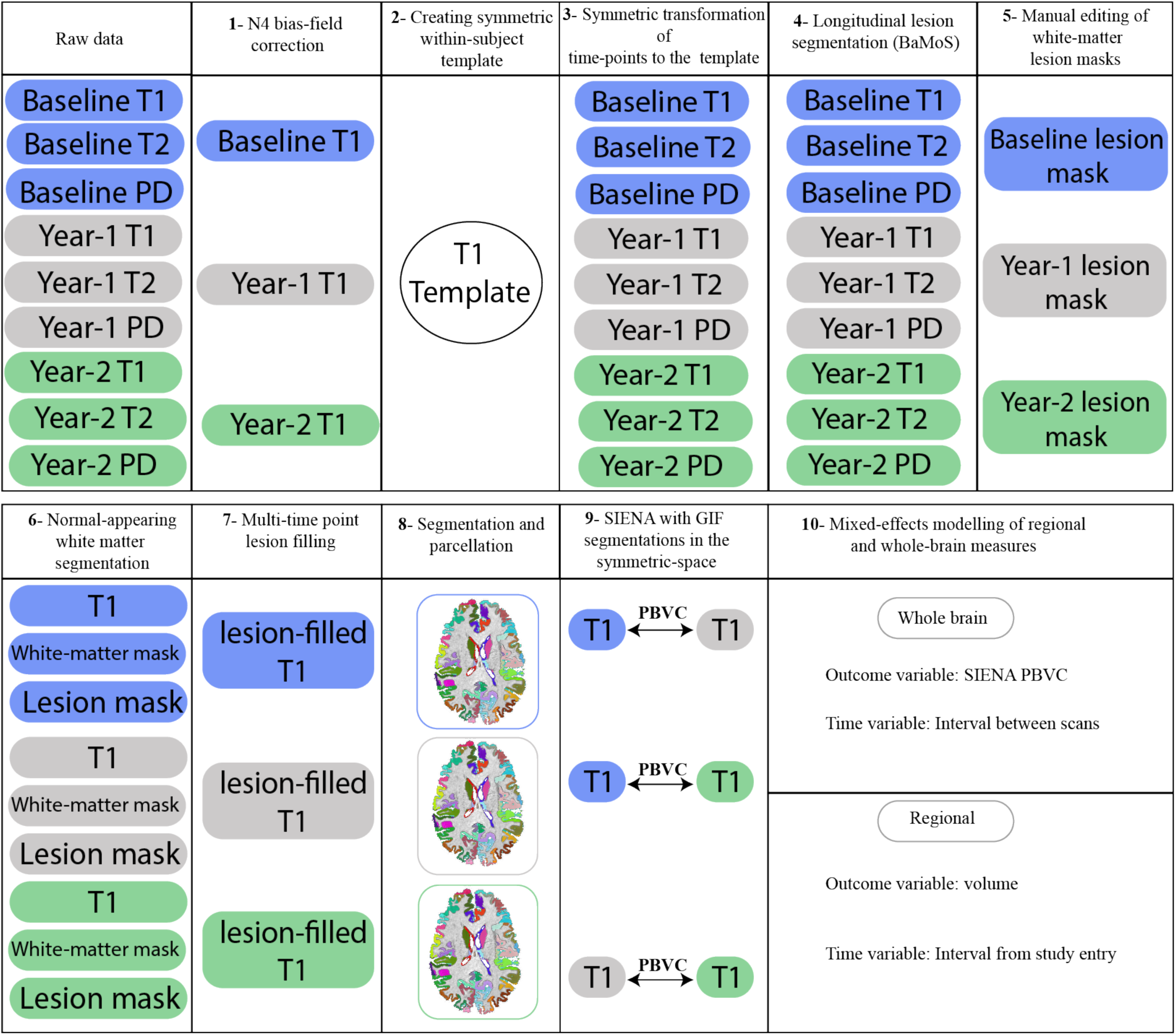
Image analysis pipeline

## Supplemental tables

**Supplementary table 1.** MRI protocol.

|  |  |  |
| --- | --- | --- |
| Sequence: T1-weighted |  |  |
| Centre | One | Two |
| Vendor | General electric Signa | Siemens |
| Magnetic field | 3 Tesla | 1.5 Tesla |
| Voxel dimension | 3D (0.93x0.93x1.1 mm) | 3D (1.25x1.25x1.2) |
| Repetition time (TR) | 7.808 ms | 2400 ms |
| Echo time (TE) | 3.004 ms | 3.45 |
| Acquisition matrix | 256x256 | 192x192 |
| Inversion time | 450 ms | 1000 ms |
| Flip angle | 20 | 8 |
| Field of view |  | 192x192 |
| Number of slices | 170 | 160 |
| Sequence: dual echo proton density and T2-weighted |  |  |
| Centre | One | Two |
| Voxel dimension | 0.97x0.97x3.0 | 0.48x0.48x3.0 |
| Acquisition type | 2D | 2D |
| Repetition time (TR) | 2600 ms | 4220 ms |
| Spacing between slices | 3 mm | 3 mm |
| Echo train length | 10 | 5 |
| Acquisition matrix | 256x256 | 424x512 |
| Flip angle | 90 | 150 |
| Number of slices | 46 | 46 |

**Supplementary Table 2.** Baseline characteristics of the participants.

|  | Treatment | Placebo | Combined |
| --- | --- | --- | --- |
| No. of participants | 67 | 64 | 131 |
| Average age in years (SD) | 51.58 (7.03) | 50.9 (6.94) | 51.2 (6.97) |
| Females (males) | 46 (21) | 42 (22) | 88 (43) |
| Duration of SPMS in years<br>(SD) | 7.41 (5.74) | 6.87 (4.62) | 7.14 (5.21) |
| Duration of MS in years (SD) | 22.32 (8.27) | 19.97 (8.94) | 21.17 (8.65) |
| Median EDSS (range) | 6 (6-6.5) | 6 (4-7) | 6 (4-7) |
| Average years of education<br>(SD) | 13.69 (3.07) | 13.42 (3.2) | 13.56 (3.16) |

*Abbreviations:* SD; standard deviation, SPMS; secondary-progressive multiple sclerosis, MS; multiple sclerosis, EDSS; Kurtzke's Expanded-Disability Status Scale.

## References

Altman DG, Bland JM. Statistics notes: Absence of evidence is not evidence of absence. BMJ 1995; 311: 485–485.

Association WM. Declaration of Helsinki, ethical principles for medical research involving human subjects. 52 Nd WMA Gen. Assem. Edinb. Scotl. 2000

Avants BB, Epstein CL, Grossman M, Gee JC. Symmetric diffeomorphic image registration with cross-correlation: evaluating automated labeling of elderly and neurodegenerative brain. Med Image Anal 2008; 12: 26–41.

Benjamini Y, Hochberg Y. Controlling the False Discovery Rate: A Practical and Powerful Approach to Multiple Testing. J. R. Stat. Soc. Ser. B Methodol. 1995; 57: 289–300.

Bollen KA, Long JS. Tests for Structural Equation Models: Introduction. Sociol. Methods Res. 1992; 21: 123–131.

Bosma LV a. E, Sonder JM, Kragt JJ, Polman CH, Uitdehaag BMJ. Detecting clinically-relevant changes in progressive multiple sclerosis. Mult. Scler. Houndmills Basingstoke Engl. 2015; 21: 171–179.

Boyes RG, Gunter JL, Frost C, Janke AL, Yeatman T, Hill DLG, et al. Intensity non-uniformity correction using N3 on 3-T scanners with multichannel phased array coils. NeuroImage 2008; 39: 1752–1762.

Carass A, Roy S, Jog A, Cuzzocreo JL, Magrath E, Gherman A, et al. Longitudinal multiple sclerosis lesion segmentation: Resource and challenge. NeuroImage 2017; 148: 77–102.

Carassiti D, Altmann DR, Petrova N, Pakkenberg B, Scaravilli F, Schmierer K. Neuronal loss, demyelination and volume change in the multiple sclerosis neocortex. Neuropathol. Appl. Neurobiol. 2017

Cardoso MJ, Modat M, Wolz R, Melbourne A, Cash D, Rueckert D, et al. Geodesic information flows: spatially-variant graphs and their application to segmentation and fusion. IEEE Trans. Med. Imaging 2015; 34: 1976–1988.

Chan D, Binks S, Nicholas JM, Frost C, Cardoso MJ, Ourselin S, et al. Effect of high-dose simvastatin on cognitive, neuropsychiatric, and health-related quality-of-life measures in secondary progressive multiple sclerosis: secondary analyses from the MS-STAT randomised, placebo-controlled trial. Lancet Neurol. 2017; 16: 591–600.

Chataway J, Schuerer N, Alsanousi A, Chan D, MacManus D, Hunter K, et al. Effect of high-dose simvastatin on brain atrophy and disability in secondary progressive multiple sclerosis (MS-STAT): a randomised, placebo-controlled, phase 2 trial. Lancet Lond. Engl. 2014; 383: 2213–2221.

Douaud G, Refsum H, de Jager CA, Jacoby R, Nichols TE, Smith SM, et al. Preventing Alzheimer’s disease-related gray matter atrophy by B-vitamin treatment. Proc. Natl. Acad. Sci. U. S. A. 2013; 110: 9523–9528.

Dubois B, Slachevsky A, Litvan I, Pillon B. The FAB: a Frontal Assessment Battery at bedside. Neurology 2000; 55: 1621–1626.

Eshaghi A, Prados F, Brownlee W, Altmann DR, Tur C, Cardoso MJ, et al. Deep grey matter volume loss drives disability worsening in multiple sclerosis [Internet]. bioRxiv 2017Available from: http://biorxiv.org/content/early/2017/08/29/182006.abstract

Eshaghi A, Prados F, Brownlee W, Altmann DR, Tur C, Cardoso MJ, et al. Deep grey matter volume loss drives disability worsening in multiple sclerosis. Ann. Neurol. 2018

Fox NC, Ridgway GR, Schott JM. Algorithms, atrophy and Alzheimer’s disease: Cautionary tales for clinical trials. NeuroImage 2011; 57: 15–18.

Frischer JM, Bramow S, Dal-Bianco A, Lucchinetti CF, Rauschka H, Schmidbauer M, et al. The relation between inflammation and neurodegeneration in multiple sclerosis brains. Brain J. Neurol. 2009; 132: 1175–1189.

Greenwood J, Mason JC. Statins and the vascular endothelial inflammatory response. Trends Immunol. 2007; 28: 88–98.

Greenwood J, Steinman L, Zamvil SS. Statin therapy and autoimmune disease: from protein prenylation to immunomodulation. Nat. Rev. Immunol. 2006; 6: 358–370.

Gronwall DMA. Paced Auditory Serial-Addition Task: A Measure of Recovery from Concussion. Percept. Mot. Skills 1977; 44: 367–373.

Groth-Marnat G, Teal M. Block design as a measure of everyday spatial ability: a study of ecological validity. Percept. Mot. Skills 2000; 90: 522–526.

Hartung JohnPD, Cottrell James EMD, Giffin Joseph P.MD Absence of Evidence Is not Evidence of Absence. Anesthesiology 1983; 58: 298–299.

Hawker K, O’Connor P, Freedman MS, Calabresi PA, Antel J, Simon J, et al. Rituximab in patients with primary progressive multiple sclerosis: results of a randomized double-blind placebo-controlled multicenter trial. Ann. Neurol. 2009; 66: 460–471.

Henneman WJP, Sluimer JD, Barnes J, van der Flier WM, Sluimer IC, Fox NC, et al. Hippocampal atrophy rates in Alzheimer disease: added value over whole brain volume measures. Neurology 2009; 72: 999–1007.

Hobart J, Lamping D, Fitzpatrick R, Riazi A, Thompson A. The Multiple Sclerosis Impact Scale (MSIS-29): a new patient-based outcome measure. Brain J. Neurol. 2001; 124: 962–973.

Hu L, Bentler PM. Cutoff criteria for fit indexes in covariance structure analysis: Conventional criteria versus new alternatives. Struct. Equ. Model. Multidiscip. J. 1999; 6: 1–55.

Imai K, Keele L, Tingley D, Yamamoto T. Unpacking the Black Box of Causality: Learning about Causal Mechanisms from Experimental and Observational Studies. Am. Polit. Sci. Rev. 2011; 105: 765–789.

Josephs KA, Dickson DW, Tosakulwong N, Weigand SD, Murray ME, Petrucelli L, et al. Rates of hippocampal atrophy and presence of post-mortem TDP-43 in patients with Alzheimer’s disease: a longitudinal retrospective study. Lancet Neurol. 2017; 16: 917–924.

Kievit RA, Davis SW, Mitchell DJ, Taylor JR, Duncan J, Team C-CR, et al. Distinct aspects of frontal lobe structure mediate age-related differences in fluid intelligence and multitasking. Nat. Commun. 2014; 5: 5658.

Klein A, Tourville J. 101 labeled brain images and a consistent human cortical labeling protocol [Internet]. Front. Neurosci. 2012; 6[cited 2016 Jul 7] Available from: http://journal.frontiersin.org/article/10.3389/fnins.2012.00171/abstract

Kurtzke JF. Rating neurologic impairment in multiple sclerosis: an expanded disability status scale (EDSS). Neurology 1983; 33: 1444–1452.

Larochelle C, Uphaus T, Prat A, Zipp F. Secondary Progression in Multiple Sclerosis: Neuronal Exhaustion or Distinct Pathology? Trends Neurosci. 2016; 39: 325–339.

Leung KK, Ridgway GR, Ourselin S, Fox NC, Alzheimer’s Disease Neuroimaging I. Consistent multi-time-point brain atrophy estimation from the boundary shift integral. Neuroimage 2012; 59: 3995–4005.

Malone IB, Leung KK, Clegg S, Barnes J, Whitwell JL, Ashburner J, et al. Accurate automatic estimation of total intracranial volume: a nuisance variable with less nuisance. NeuroImage 2015; 104: 366–372.

Merkle EC, Rosseel Y. blavaan: Bayesian structural equation models via parameter expansion [Internet]. ArXiv151105604 Stat 2015[cited 2018 Mar 1] Available from: http://arxiv.org/abs/1511.05604

Montalban X, Hauser SL, Kappos L, Arnold DL, Bar-Or A, Comi G, et al. Ocrelizumab versus Placebo in Primary Progressive Multiple Sclerosis. N. Engl. J. Med. 2017; 376: 209–220.

Pinheiro J, Bates D, DebRoy S, Sarkar D, R Core Team. nlme: Linear and Nonlinear Mixed Effects Models [Internet]. 2017[cited 2017 Oct 18] Available from: https://cran.rproject.org/web/packages/nlme/citation.html

Polman CH, Reingold SC, Edan G, Filippi M, Hartung HP, Kappos L, et al. Diagnostic criteria for multiple sclerosis: 2005 revisions to the ‘McDonald Criteria’. Ann Neurol 2005; 58: 840–6.

Prados F, Cardoso MJ, Kanber B, Ciccarelli O, Kapoor R, Gandini Wheeler-Kingshott CAM, et al. A multi-time-point modality-agnostic patch-based method for lesion filling in multiple sclerosis. NeuroImage 2016; 139: 376–384.

R Core Team. R: A Language and Environment for Statistical Computing [Internet]. Vienna, Austria: R Foundation for Statistical Computing; 2014. Available from: http://www.Rproject.org/

Reuter M, Fischl B. Avoiding asymmetry-induced bias in longitudinal image processing. NeuroImage 2011; 57: 19–21.

Rosseel Y. lavaan: An *R* Package for Structural Equation Modeling [Internet]. J. Stat. Softw. 2012; 48[cited 2017 Aug 26] Available from: http://www.jstatsoft.org/v48/i02/

Smith SM. Fast robust automated brain extraction. Hum Brain Mapp 2002; 17: 143–55.

Smith SM, De Stefano N, Jenkinson M, Matthews PM. Normalized accurate measurement of longitudinal brain change. J. Comput. Assist. Tomogr. 2001; 25: 466–475.

Smith SM, Rao A, De Stefano N, Jenkinson M, Schott JM, Matthews PM, et al. Longitudinal and cross-sectional analysis of atrophy in Alzheimer’s disease: Cross-validation of BSI, SIENA and SIENAX. NeuroImage 2007; 36: 1200–1206.

Sudre CH, Cardoso MJ, Bouvy WH, Biessels GJ, Barnes J, Ourselin S. Bayesian model selection for pathological neuroimaging data applied to white matter lesion segmentation. IEEE Trans. Med. Imaging 2015; 34: 2079–2102.

Sudre CH, Cardoso MJ, Ourselin S, Alzheimer’s Disease Neuroimaging Initiative. Longitudinal segmentation of age-related white matter hyperintensities. Med. Image Anal. 2017; 38: 50–64.

Thompson AJ, Baranzini SE, Geurts J, Hemmer B, Ciccarelli O. Multiple sclerosis. The Lancet 2018; 391: 1622–1636.

Tustison NJ, Avants BB, Cook PA, Zheng Y, Egan A, Yushkevich PA, et al. N4ITK: improved N3 bias correction. IEEE Trans. Med. Imaging 2010; 29: 1310–1320.

Wagenmakers E-J, Farrell S. AIC model selection using Akaike weights. Psychon. Bull. Rev. 2004; 11: 192–196.

Wechsler D. Wechsler Abbreviated Scale of Intelligence. 2011

Weitz-Schmidt G, Welzenbach K, Brinkmann V, Kamata T, Kallen J, Bruns C, et al. Statins selectively inhibit leukocyte function antigen-1 by binding to a novel regulatory integrin site. Nat. Med. 2001; 7: 687–692.

Zhang Y, Brady M, Smith S. Segmentation of brain MR images through a hidden Markov random field model and the expectation-maximization algorithm. IEEE Trans Med Imaging 2001; 20: 45–57.

